# Dendritic morphology constrains microtubule dynamics and organization

**DOI:** 10.64898/2026.09.28.755114

**Authors:** Dongdong You, Louison Thorens, Zhi Xu, Yun Fu, Ning Wang, Timothy J. Mitchison, Maijia Liao

**Affiliations:** Department of Physics, Northeastern University, Boston, MA, USA; Department of Bioengineering, Northeastern University, Boston, MA, USA; Institute for Mechanobiology, Northeastern University, Boston, MA, USA; Department of ECE and Khoury College of Computer Science, Northeastern University, Boston, MA, USA; Department of Systems Biology, Harvard Medical School, Boston, MA, USA

**Author notes:** These authors contributed equally to this work (Dongdong You, Louison Thorens).

## Abstract

Neuronal dendrites form highly branched morphologies supported by spatially patterned microtubule arrays that are essential for intracellular transport and structural stability, yet how this organization is established and maintained remains unclear. Here we analyze microtubule behavior in *Drosophila* sensory neurons using live-cell imaging, deep learning–based analysis, and genetic perturbations. We find that microtubules are shorter and fewer in thin distal branches and progressively longer and more abundant in thicker proximal branches. Locally, polymerization slows in narrow branches due to soluble tubulin depletion, while catastrophe frequency increases near branch points upon cortical encounter. A physical model incorporating these morphology-dependent constraints reproduces both local microtubule dynamics and global organization. Inverting the textbook view that microtubule dynamics shape cell morphology, our results demonstrate that, on short timescales, cell morphology instead constrains microtubule dynamics, thereby determining large-scale cytoskeletal organization.

## Introduction

Neuronal dendrites exhibit elaborate branching morphologies that remodel and ramify as the innervated tissues grow (1, 2). Dendrite remodeling and homeostasis require a precisely organized microtubule cytoskeleton that shapes dendritic architecture and serves as ‘rails’ for the transport of mRNAs, organelles, and protein cargos such as signaling complexes (3, 4). While dendritic remodeling occurs over the course of days, individual microtubules undergo dynamic instability, alternating between polymerization and depolymerization within seconds to minutes (5–8). How stochastic dynamic microtubules self-organize into ordered arrays within slowly remodeling and highly branched dendrites, and thereby maintain dendritic homeostasis, remains poorly understood.

Over the past decades, individual microtubule dynamics have been extensively studied in diverse systems, including reconstituted *in vitro* assays, *C. elegans*, Xenopus egg extracts, plants, and cultured cells (8–14). While informative, these studies largely focus on systems with simplified cellular architecture and do not capture the complexity of highly branched dendrites. Individual microtubules within neuronal compartments can be resolved by electron microscopy reconstructions (15, 16), but these approaches cannot reveal their dynamics. Moreover, the restricted spatial extent of such reconstructions is insufficient to determine microtubule organization across large and complex dendritic arbors. Complementing these studies, work in reconstituted systems and living cells have shown that cellular geometry can influence large-scale cytoskeletal organization (17–20). However, the local dynamic mechanisms through which complex cellular geometries regulate microtubule behavior to generate this organization remain unknown. Together, these limitations highlight the need for direct measurements that link microtubule dynamics to higher-order organization in highly branched dendrites within intact nervous systems.

To address this gap, we turned to *Drosophila* class IV dendritic arborization neurons, which form a dense, highly branched meshwork beneath the cuticle and are amenable to *in vivo* imaging in whole animals (21–24). These neurons detect noxious stimuli, including heat (25), ultra-violet light (26), and harsh mechanical stimuli (27) over the larval surface (Fig. 1A). The systematic tapering of dendritic diameters from primary branches toward periphery is captured by our previously defined metric, the leaf number (28), which counts the number of terminal branches supported by each branch and decreases toward the periphery (Fig. 1B). This hierarchical organization generates dendritic arbors composed of interconnected compartments spanning a wide range of volumes in which microtubules reside. Collectively, these features make class IV sensory neurons a canonical system for studying how microtubules adapt to complicated cellular morphology.

**Figure 1.**
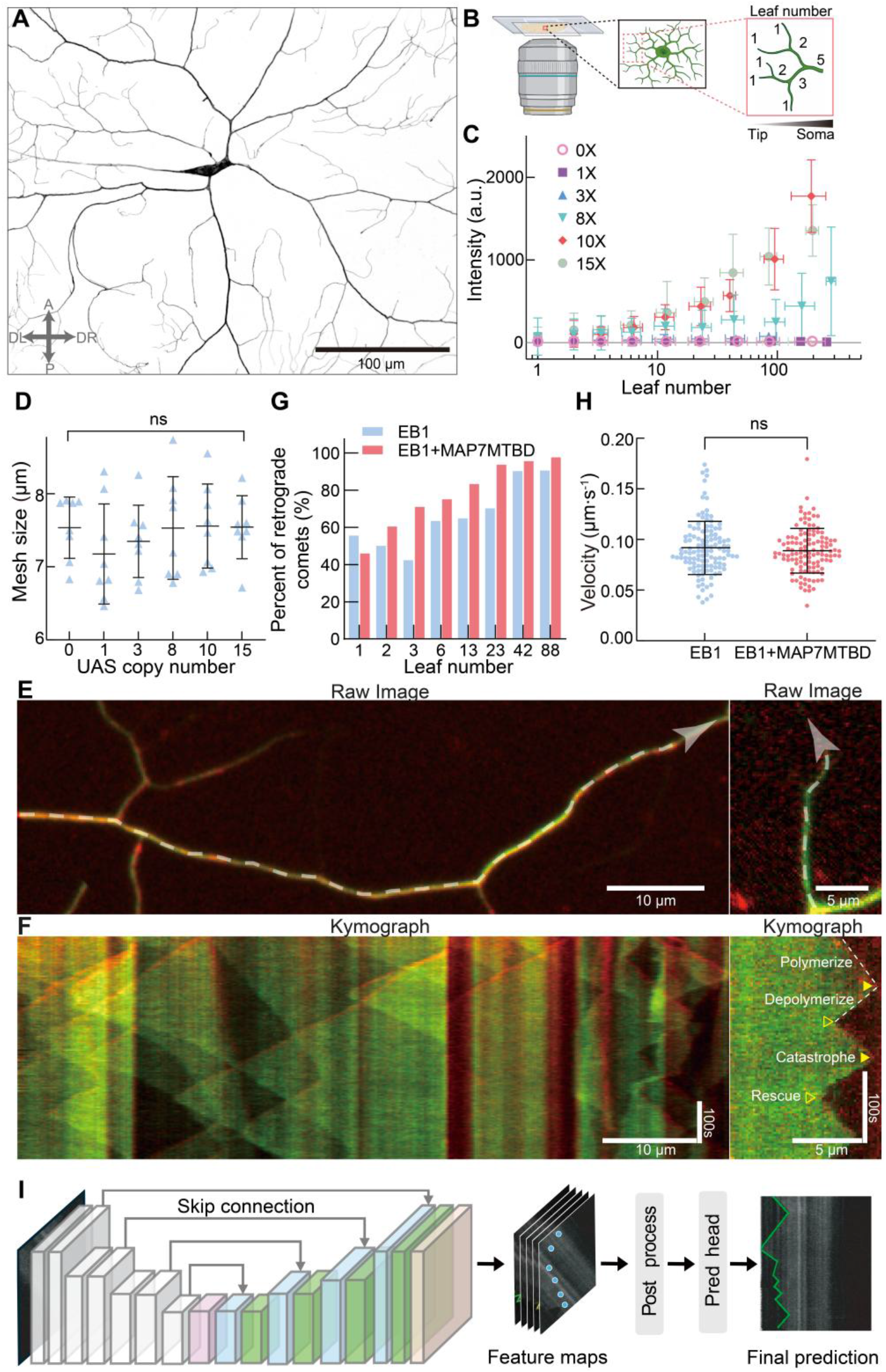
MAP7MTBD enables *in vivo* visualization of microtubule lattice and dynamics without altering morphology in *Drosophila* sensory neurons. (A) Confocal image of a Class IV neuron with ppkCD4-tdTomato. (B) Schematic illustrating the imaging setup and leaf number definition. The gray triangle indicates increasing leaf number toward the soma. (C) Average MAP7MTBD-mNeonGreen intensity along branches as a function of leaf number for different UAS copy numbers (three neurons per condition, points indicate mean, error bars denote SD). (D) Mesh size is unchanged across UAS copy numbers (Materials and Methods, p = 0.70, Brown–Forsythe ANOVA, N = 8 animals for each copy number). Black lines show mean ± SD. (E–F) Confocal images and corresponding kymographs of primary and peripheral dendrites co-labeled with EB1-RFP and MAP7MTBD-mNeonGreen. White dashed curves indicate the kymograph measurement paths; arrows indicate direction. Microtubule dynamics events: catastrophe, rescue, polymerization, and depolymerization are shown in F-right. (G) Fraction of retrograde EB1 comets as a function of leaf number. (H) EB1 comet velocities with and without MAP7MTBD show no significant difference (p = 0.62, Welch’s t test, N = 3 animals). Black lines indicate mean ± SD. (I) Illustration of the KymoNet workflow for automated microtubule dynamics detection.

Leveraging this system, we combined direct visualization with deep learning and analytical approaches to quantify both microtubule dynamics and organizational features. By integrating these measurements with computational modeling, we revealed that neuronal branching morphology and tubulin abundance jointly regulate microtubule dynamics, giving rise to higher-order microtubule organization that scales with branch volume. Our findings demonstrate that dendrite morphology is sufficient to organize microtubules and thereby support dendritic homeostasis on short timescales, in contrast to the usual assumption that microtubules determine cell morphology.

## Results

### Dynamic microtubules can be visualized in *Drosophila* neuronal dendrites

Direct visualization of microtubule lattice dynamics in *Drosophila* sensory neurons *in vivo* remains notoriously difficult, as tagging tubulin dimers with bulky fluorophores can perturb their dynamics (29). To overcome this limitation, we expressed the microtubule-binding domain of MAP7 (MAP7MTBD), which decorates the entire microtubule lattice without perturbing polymerization dynamics or transport (30–34). We fused MAP7MTBD to mNeonGreen (35) and systematically varied its expression levels by adjusting UAS copy numbers (Fig. 1C). Dendritic morphology remained unchanged across this range, confirming no perturbation even at high expression levels (Fig. 1D). We selected an intermediate level of expression that provided optimal visualization of microtubule dynamics. Using fluorescence recovery after photobleaching (FRAP), we found that ~92% of MAP7MTBD was lattice-bound, which explains the high contrast of microtubule labeling. To validate our scoring of individual microtubules undergoing dynamic instability we co-expressed EB1-RFP as a plus-end marker. The polymerization phase is clearly marked by EB1 comets (36–38), whereas the onset of depolymerization closely followed the disappearance of these comets (Figs. 1E-F). EB1 velocity and microtubule polarity (Figs. 1G-H) remained consistent over a large range of MAP7MTBD expression levels, indicating that our labeling strategy preserves native microtubule behavior.

### Quantification of microtubule dynamics

We developed a deep-learning-based analysis pipeline, KymoNet (Fig. 1I), to collect statistically sufficient data across a full range of neuronal dendrite leaf numbers and we quantified the four dynamic parameters describing microtubules dynamic instability: polymerization rate *v*_*g*_, depolymerization rate *v*_*s*_, catastrophe frequency *f*_*c*_, and rescue frequency *f*_*r*_ (Fig. 1F). Since the same cytoplasm is shared between dendrites, we expect these parameters to be spatially invariant. Surprisingly, three of the four dynamic parameters were significantly correlated with leaf number (Figs. 2A-B). Both polymerization and depolymerization rates increased with increasing leaf number before they plateaued, whereas the catastrophe frequency decreased with leaf number and reached a plateau near the soma. Rescue frequency, however, remained largely constant irrespective of the branch order.

**Figure 2.**
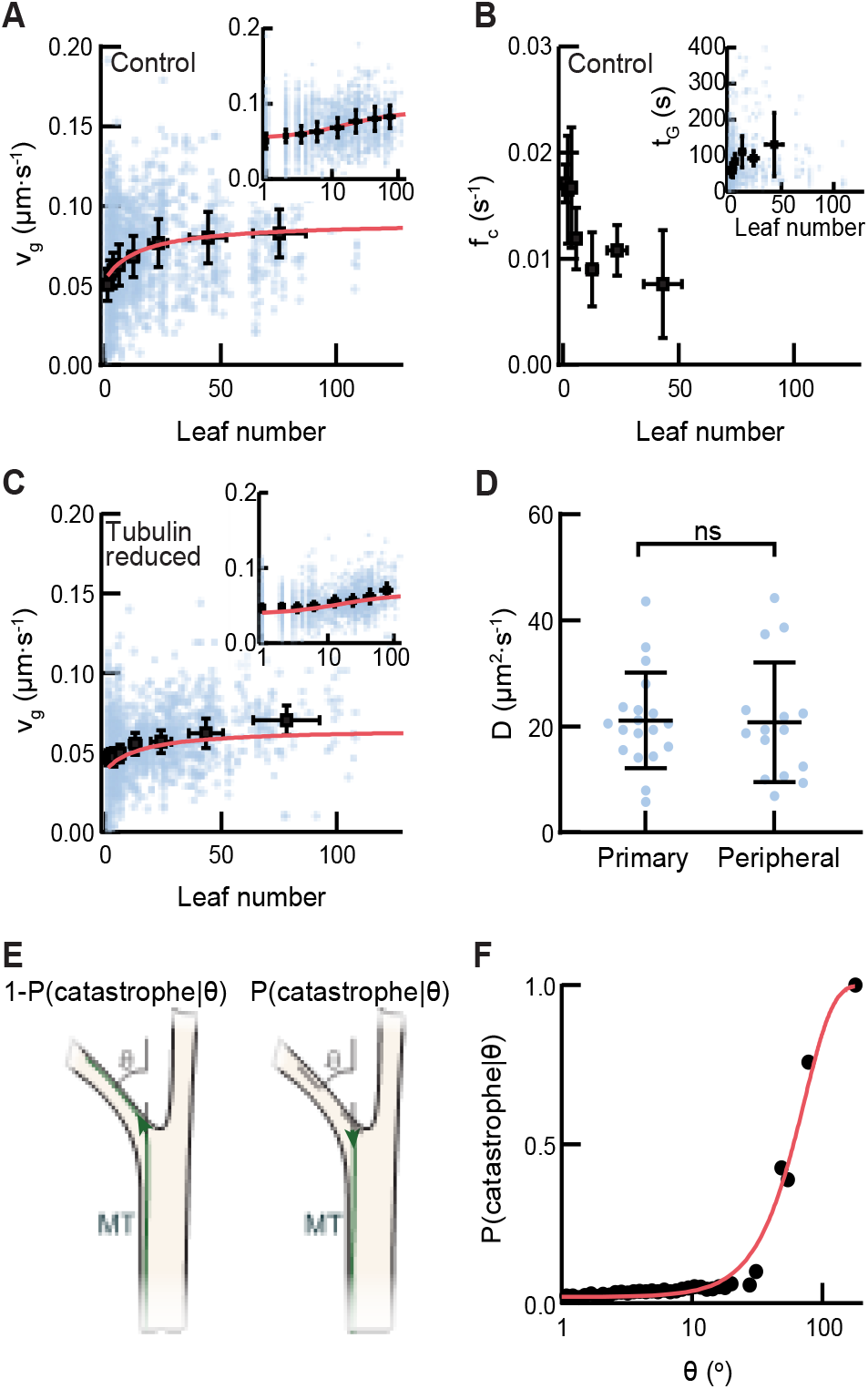
Microtubule dynamics characterization in dendritic geometry. (A) Polymerization rate *v*_*g*_, (B) catastrophe frequency *f*_*c*_ in control neurons (N = 6) as a function of leaf number. (C) *v*_*g*_ in tubulin-reduced neurons (N = 7) as a function of leaf number. Blue dots show all individual measurements; black squares denote means within logarithmic leaf-number bins (error bars: standard deviation within each bin). Insets in (A, C) show the same data on a log-scaled x-axis to highlight low-leaf-number behavior. The inset in (B) shows the average growth-time vs. leaf number used to estimate local catastrophe frequencies. Pearson coefficients and p values for the log-transformed leaf number: (A) 0.99 (p < 0.001), (B) −0.92 (p = 0.003), (C) 0.94 (p < 0.001). Fits in (A, C) used Eq. (1) (red curve) with *c*_0_ = 8.64 ± 0.80 μM (mean ± SE) and *c*_0_ = 6.25 ± 0.92 μM (mean ± SE) respectively. (D) Free-tubulin diffusion coefficient in primary and peripheral (leaf number < 10) branches; 3–4 measurements per neuron from six animals. (E) Schematic showing how a microtubule reacts as it encounters a local deflection angle *θ*. (F) Probability that a microtubule plus end undergoes catastrophe as a function of the deflection angle *θ*.

### Dendritic diameter regulates polymerization rate by regulating soluble tubulin concentration

Microtubule polymerization rates depend on the concentration of free tubulin subunits (9). To test whether this relationship holds *in vivo*, we reduced the tubulin level by using heterozygous *aTub84B*^*attp-KO*^ mutant flies and found that the microtubule polymerization rate decreased across all leaf numbers (Fig. 2C). These results indicate that the observed attenuation of polymerization rate toward peripheral branches with smaller caliber reflects a local decrease in free tubulin concentration. We therefore sought to quantitatively link soluble tubulin-dependent microtubule polymerization rate to local cross-sectional area, which varies over an order of magnitude from 0.03 to 0.3 μm^2^.

Spatial regulation of soluble tubulin concentration implies that polymerization must deplete tubulin faster than mixing by diffusion in peripheral branches. We measured the diffusion coefficient of freely mobile cytoplasmic tubulin (Fig. 2D) and found it was spatially uniform, with an estimated value of *D* = 20 ± 10 μm^2^/s. This value is comparable to reported *in vivo* measurements of 6-10 μm^2^/s (39–41) and is consistent with estimates based on a cytoplasmic viscosity of ~2.7 cP, as expected for neuronal cytoplasm (42) (Methods). This diffusion coefficient, combined with microtubule polymerization rates *v*_*g*_, defines a length over which tubulin dimers diffuse before being consumed by the tip of a growing microtubule, 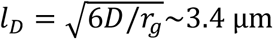, where *r*_*g*_ = *v*_*g*_/*δ* is the rate of attachment of a tubulin dimer at the tip of a microtubule and *δ* is the tubulin dimer length (43, 44). The branch diameter is significantly smaller than the diffusion length, such that diffusion rapidly equilibrates tubulin across the cross-section. We therefore approximate transport along the branch as effectively one-dimensional and build a phenomenological theory based on a drift-diffusion equation. As microtubules grow, they locally deplete free tubulin, and this depletion becomes more pronounced as branch radius narrows, as predicted by equation (1):

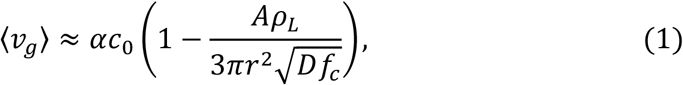

where *ρ*_*L*_ is the linear density of tubulin dimmers, *r* is the branch radius, *c*_0_ is the free tubulin concentration, and *A* is a prefactor. This theory nicely accounts for polymerization rate variations in both control and tubulin-reduced neurons (Figs. 2A-C) and gives a direct readout of the free tubulin concentration *c*_0_. We thus conclude that reduced branch cross-sectional area is sufficient to account for the decreased polymerization rate in peripheral branches.

### Local branching orientation regulates microtubule catastrophe

Next, we asked what factors govern microtubule catastrophe. Soluble tubulin concentration regulates catastrophe rates in some models (9, 45) but not others (5, 46–48). We observed no significant change in catastrophe rate upon altering tubulin concentration. Thus, we sought alternative explanations for the spatial regulation of catastrophe.

Constraints imposed by cellular morphology, including the cell cortex, are known to promote catastrophes (49–52). When a growing microtubule encounters a geometric constraint, it can either reorient to a new path or undergo catastrophe and begin depolymerization (53). We therefore asked whether branch morphology modulates catastrophe. Because microtubules are confined within narrow dendritic calibers, their deflection angle can be approximated by the local change in branch orientation *θ* (Fig. 2E). The probability of catastrophe following deflection, *P*(catastrophe|*θ*), increased monotonically with *θ* under both control and reduced tubulin levels (Fig. 2F). Sensory neuronal dendrites are highly curved, with peripheral branch points exhibiting sharper deflection, implying correspondingly higher catastrophe rates.

### Microtubules are shorter and less abundant in peripheral branches

The leaf-number dependence of microtubule dynamics motivated us to examine microtubule organization along dendritic arbors. Since microtubules are densely packed in dendrites, their length and number have been estimated using electron microscopy reconstructions (15, 16, 54, 55) or expansion microscopy approaches (56) that are labor-intensive and restricted to fixed samples (57). Although diffraction-limited light microscopy cannot resolve individual microtubules, our ability to track their dynamics enabled a kymograph-based approach (10) to extract single microtubule intensity. After subtracting the cytoplasmic MAP7MTBD fraction to isolate the lattice-bound component, we used the single microtubule intensity as a reference to map microtubule distribution along each branch (Fig. 3A). Stepwise increases and decreases marked microtubule start and end sites, allowing simultaneous measurement of microtubule length and number. We further validated this approach using synthetic images generated from experimental parameters and found it reliable up to 55 microtubules (<15% relative error, Fig. 3B), exceeding the maximal microtubule counts we observed.

**Figure 3.**
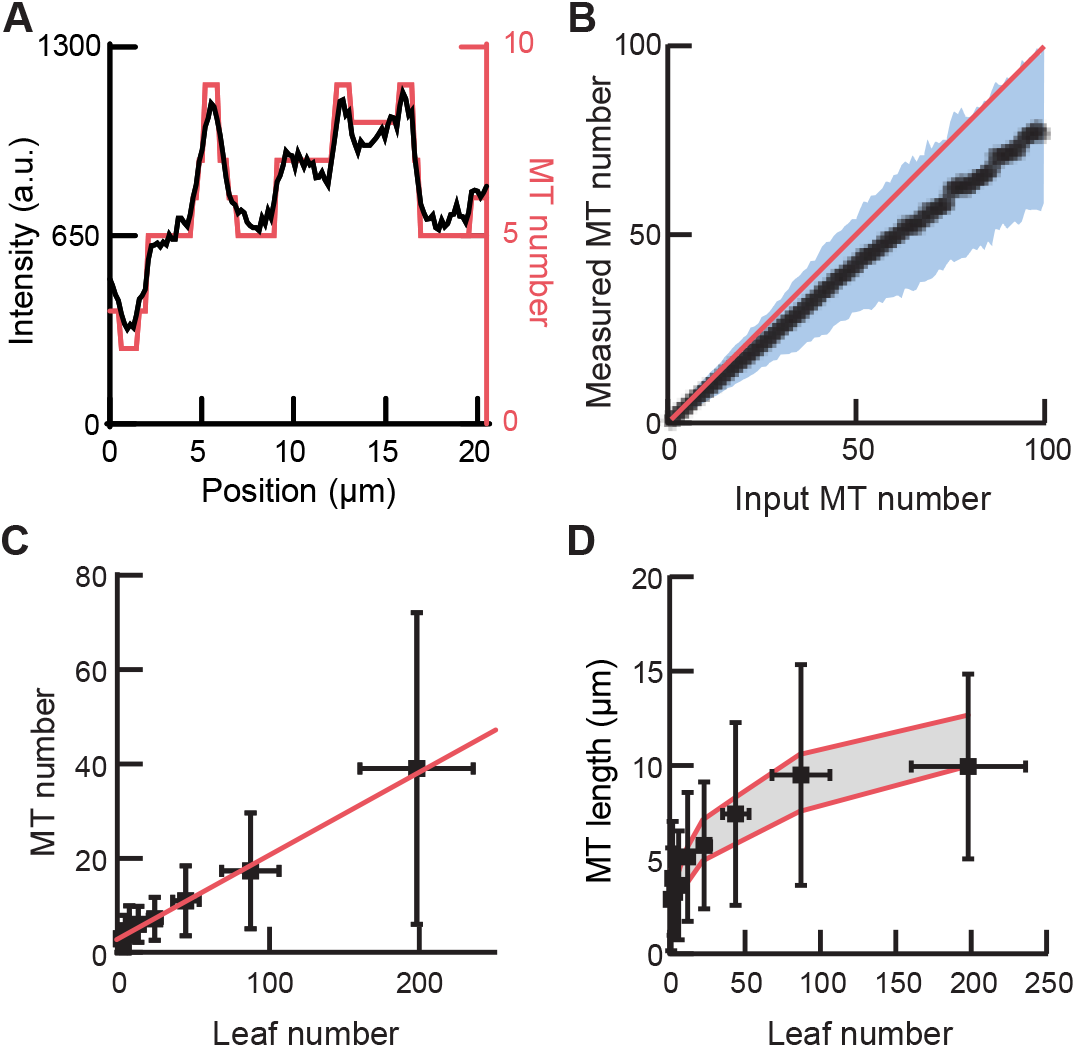
Microtubule distribution across dendritic arbors. (A) Representative microtubule intensity along a branch after background subtraction (black, left axis). The quantized microtubule number distribution is shown in red (right axis). (B) Validation of total microtubule number quantification using synthetic images. The blue region indicates the standard deviation (N = 5000 simulated microtubules per ground-truth value). The red line denotes y = x. (C) Total microtubule number as a function of leaf number. Error bars indicate standard deviation across branches. The red line shows a linear fit with slope 0.18. (D) Average microtubule length as a function of leaf number. Because microtubule dynamic parameters are consistent within branches of the same leaf number, leaf number is used to define branch identity. Experimental measurements are compared with theoretical predictions assuming microtubules are measured within branches of length *L*_*i*_ for each leaf number. These boundary conditions constrain the average microtubule length to lie between a lower bound, ⟨*l*⟩ = ⟨*λ* − *L*_*i*_ exp(−*L*_*i*_/*λ*) /(1 − exp(−*L*_*i*_/*λ*))⟩_*i*_ corresponding to a hard wall boundary, and an upper bound, ⟨*l*⟩ = ⟨*λ*(1 − exp(−*L*_*i*_/*λ*))⟩_*i*_ corresponding to an open boundary, where *λ* is the microtubule characteristic length. Error bars indicate standard deviation.

We found microtubule number scaled linearly with the leaf number with a slope of 0.18 (Fig. 3C). From our previously established relation between leaf number and branch cross-sectional area (28), we inferred an average nearest-neighbor microtubule spacing of ~100 nm, consistent with electron microscopy measurements (58, 59). The average microtubule length increased monotonically from 2.9 ± 2.7 μm in terminal branches to 9.5 ± 5.9 μm in primary branches (mean ± SD, Fig. 3D), consistent with an independent measurement of single microtubule length. To understand what sets the length, we turned to a two-state model of dynamic instability that connects steady-state length to four dynamic parameters (60), with an additional constraint imposed by the finite branch length *L*, since measurements are restricted to individual branches with a given leaf number. The model shows excellent agreement with experimentally measured microtubule length (Fig. 3D). Thus, microtubule number follows branch caliber, and microtubule length follows branch length, together shaping a coherent cytoskeletal organization across the arbor.

### Mechanisms of microtubule regulation in dendrites

Next, we developed a stochastic simulation based on the minimal set of elements needed to capture the experimentally measured features, consisting of microtubule nucleation and dynamics. Microtubule translocation contributes only a minor fraction of total microtubules, indicating that active microtubule transport is negligible and the distribution microtubule minus end provides a reasonable approximation of microtubule nucleation sites (61). We therefore inferred the minus-end probability distribution from measured microtubule organization profiles and polarity. Our analysis revealed that more than 50% of microtubule minus ends are concentrated at branch points. Notably, models assuming a uniform minus-end distribution failed to capture the measured length distribution. This inferred minus-end distribution is supported by prior work, which shows enrichment of the *γ* tubulin ring complex at branch points (62, 63), where it has been proposed to function as a microtubule nucleation hotspot (37). Thus, we modeled nucleation as occurring predominantly at branch points, with new microtubules introduced at a defined nucleation rate in the simulations.

With microtubule nucleation specified, we considered how microtubule dynamics should be represented in simulations. Microtubule minus ends are typically capped by associated proteins (5) and are largely non-dynamic (64). The infrequent minus-end dynamics we observed exhibited polymerization and depolymerization rates of 0.020 μm/s and 0.040 μm/s, respectively, which are significantly slower than plus-end dynamics (65). We therefore focused exclusively on plus-end behaviors and modeled the mechanisms governing them. Polymerization rate was computed from the local tubulin concentration using equation (1), catastrophe was implemented via the measured angle-dependent probability *P*(catastrophe|*θ*) (Fig. 2F), and depolymerization rate and rescue frequency were taken from experimental measurements.

We then incorporated realistic neuronal morphology into the simulation, including local curvature and dendritic diameters (28). We tuned the nucleation rate, which is the only free parameter in the simulation, while constraining all the other parameters within the experimentally measured ranges. Microtubules self-organized into a steady state (66, 67), in which polymerization consumes the available tubulin pool and depolymerization replenishes it. A nucleation rate of 0.016 s^−1^ yielded excellent agreement with the experimental measurements (Fig. 4A). Altogether, these findings show that dendritic morphology, localized nucleation, and intrinsic microtubule dynamic rules are sufficient to generate the emergent dendritic microtubule organization.

**Figure 4.**
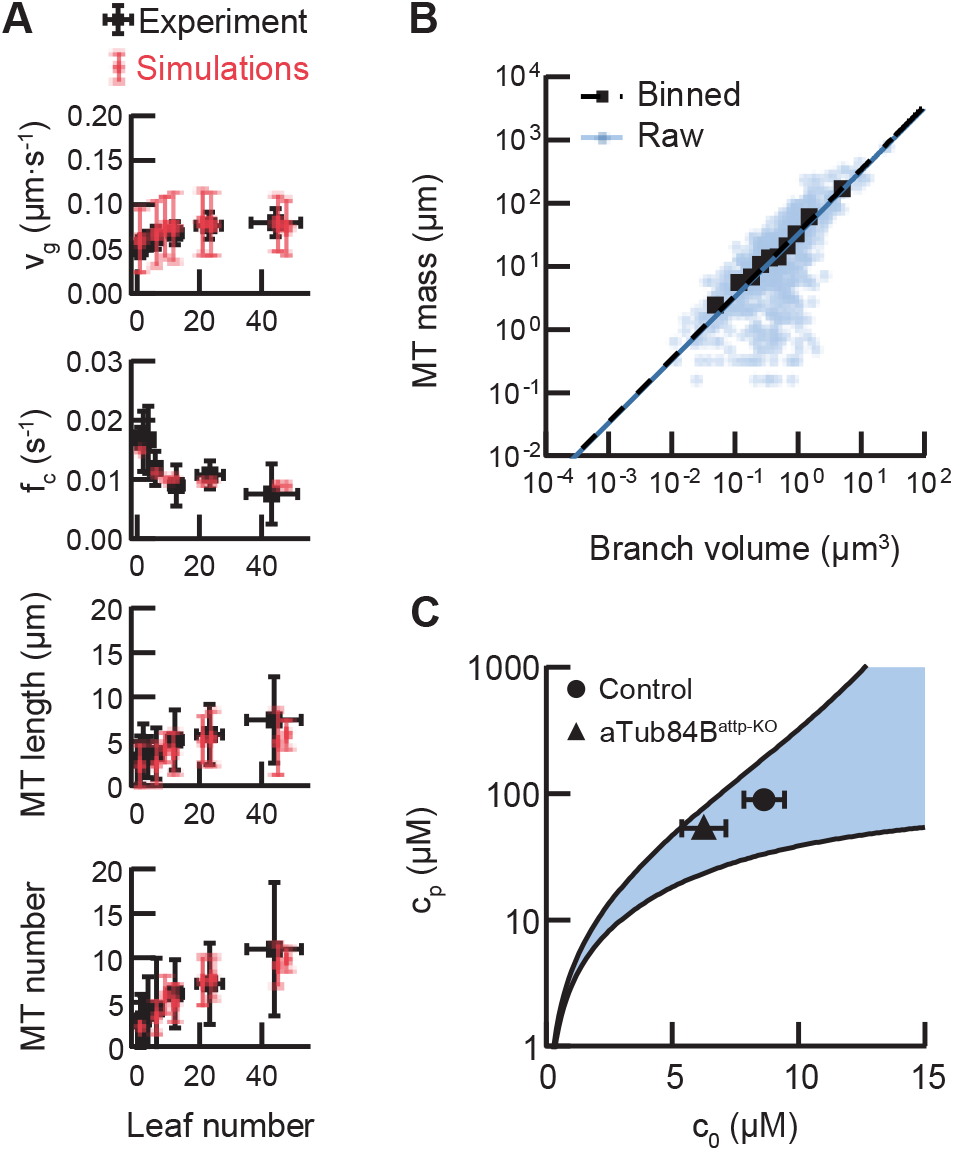
Microtubule organization is captured by simulations and tubulin partitioning. (A) Simulations results for control neurons using the optimized parameters (*f*_*N*_ = 0.016 s^−1^; *c*_total_ = 110 μM). From top to bottom: polymerization rate, catastrophe frequency, average microtubule (MT) length, and total MT number as a function of leaf number. Black squares: experiments; red circles: simulations; error bars: standard deviation. Results represent the average over 400 simulations. (B) Total microtubule mass (total microtubule length) versus branch volume for control neurons (blue circle; N = 6). To account for the uneven distribution of branch volume, data were additionally binned on a log-normal scale with equal probability bins. The binned means are shown as black squares. Solid lines represent linear fits constrained through the origin with slopes: 32.4 μm ⋅ μm^−3^ (raw data, blue), and 34.4 μm ⋅ μm^−3^ (binned data, black). The slope gives the polymerized-tubulin concentration *c*_*p*_. The value from raw data is used in the main text. (C) The measured *c*_*p*_ from panel B is plotted together with the free-tubulin concentration inferred from the polymerization-rate fits (Fig. 2A). The shaded region shows theoretical bounds from a mass balance model. Both control and reduced tubulin data fall within these bounds.

### Partitioning of tubulin in dendrites

Having established how microtubules are organized, we next asked how neurons partition tubulin between polymerized and free pools (68, 69). We found that total polymer mass scaled with branch volume, corresponding to a polymerized tubulin concentration *c*_*p*_ of 90 μM (Fig. 4B). Combined with the free tubulin concentration *c*_0_ estimated from equation (1), this yields a total tubulin concentration of 98.4 μM in control neurons, in close agreement with an independent measurement (114.5 μM). Applying the same analysis to tubulin-reduced neurons, we found that total tubulin concentration decreased by 33% relative to controls, comparable to the ~20% reduction by Western blot analysis.

To relate the cellular-scale measurements of *c*_*p*_ and *c*_0_ to microscopic details, we applied a mass-balance framework (70) based on conservation of tubulin between free and polymerized pools. At steady state, the rate of tubulin incorporation into microtubules equals the rate of tubulin loss from microtubules:

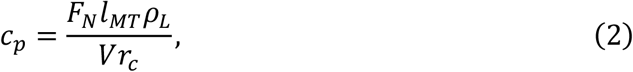

where *V* is the total volume, *r*_*c*_ is the critical nucleation rate setting the average lifetime of a microtubule, and *F*_*N*_ is the total nucleation rate determined from simulations. The effective microtubule length *l*_*MT*_ depends on local boundary conditions, reflecting whether microtubules terminate at branch points or extend into adjacent branches. Since both *l*_*MT*_ and *r*_*c*_ depend on *c*_0_, the mass-balance provides a direct link between *c*_*p*_ and *c*_0_. The experimentally measured *c*_*p*_ and *c*_0_ under both control and reduced tubulin conditions fell within the predicted bounds (Fig. 4C). Combined, our findings show microscopic rules shape tubulin partitioning at the cellular level.

## Discussion

In this work we establish a framework for directly visualizing microtubules in living sensory neurons and an analysis pipeline that quantifies microtubule dynamics and organization. By integrating these measurements with simulations, we show that macroscopic microtubule organization emerges from local dynamic rules regulating microtubule polymerization, catastrophe, and nucleation acting within the constraints of neuronal morphologies. Mass balance further explains how tubulin partitions between free and polymerized pools.

We identify that microtubule polymerization rate is determined by the balance between tubulin supply and consumption, with the predicted free tubulin levels matching independent measurements. While depolymerization rates vary spatially and independently of tubulin concentration with unclear mechanism. In primary branches, microtubules grow faster and undergo fewer catastrophes, producing longer microtubules. We hypothesize that kinesin-8 (71), a highly processive motor, may accumulate on longer microtubules and contribute to elevated depolymerization in primary branches, though this requires further study.

More broadly, our results reveal a size-dependent scaling of tubulin mass encoded by neuronal branching morphology. This scaling arises naturally from a limiting pool mechanism (17, 18, 72, 73). Smaller branches, with proportionally smaller pools, deplete subunits more rapidly during growth, slowing polymerization and causing filaments to plateau at a smaller size. When diffusion is rapid enough to equilibrate local fluctuations, the free tubulin concentration provides an indirect readout of the polymer mass, as reflected in the partitioning curve (Fig. 4C). The system is thus inherently self-correcting, as deviations from the steady state automatically shift the balance between assembly and disassembly, driving the system back toward equilibrium. This mechanism supports an emerging paradigm in which cytoskeletal organization is not governed solely by molecular regulators but is inherently encoded by cellular geometry (20, 74–76).

In this manner microtubule organization and dynamics may confer functional advantages. We find that dendritic microtubules form overlapping segments rather than continuous polymers spanning entire branches (Fig. 3A). Such a discontinuous architecture could provide local mechanical support (77) while permitting rapid adjustments as larvae crawl and dendrites deform, long continuous buckles by contrast. In addition, microtubule persistence time, defined as the mean interval between switches of polymerization and depolymerization states (78), was the shortest in terminal branches. Although this rapid remodeling comes at the cost of efficiency, it enables structural adaptability, consistent with the view that terminal branches act as a “branching engine … that initiates, directs, and maintains branch outgrowth during development and regrowth” (79, 80).

Together, our observations indicate that neuronal morphology constrains microtubule self-organization through local regulation of polymerization dynamics, thereby maintaining cytoskeletal homeostasis on short timescales over which cellular morphology remains effectively unchanged. Microtubules are known to reciprocally shape cell morphology, either by actively shaping cellular form (81, 82), or, in neurons, by directing intracellular transport and spatially organizing signaling at the cortex (3, 83, 84). These processes likely form a coupled feedback loop in which morphology determines microtubule organization at short timescales, while microtubules, in turn, drive the evolution of neuronal form at longer timescales, giving rise to a self-organizing system operating across multiple timescales (85). Our discovery therefore inverts the textbook view wherein microtubules are proposed to determine cell geometry and calls for future modeling approaches that encompass multiple scales of time.

## Materials and Methods

### Fly strains

#### The following *Drosophila* melanogaster stocks were used in this study

*w*; Sp/CyO; Dr/TM6B, w*; ppk-Gal4; Dr/TM6B, w*; Sp/CyO; ppk-Gal4, w*; ppk-Gal4; ppk-CD4-tdTomato, w*; Sp/CyO; UAS-CD4-IFP/TM6B* (gift from Chun Han’s lab), *w*; +/+; αTub84B*^*attp-KO*^*/TM3Sb kr>GFP*, and *w*; Sp/CyO; UAS-EB1-RFP/TM6B* (gift from Melissa Rolls’s Lab).

#### Stocks generated in this study included

*w*; 10XUAS-mNeonGreen/Cyo; Dr/Tm6B, w*; ppk-Gal4; ppk-Gal4::UAS-EB1-RFP::10XUAS-MAP7MTBD-mNeonGreen::UAS-CD4-IFP/TM6B, w*; Sp/Cyo; ppk-gal4::UAS-EB1-RFP::10XUASMAP7MTBD-mNeonGreen/Tm6B* and the *UAS-MAP7MTBD-mNeonGreen* series (1X, 3X, 8X, 10X, and 15X), all maintained over *Sp/CyO; TM6B*.

### Plasmid construction and generation of transgenic flies

To generate the 10XUAS-IVS-Syn21-mNeonGreen-p10 and 10XUAS-IVS-Syn21-MAP7MTBD-mNeonGreen-p10 constructs, we synthesized both the mNeonGreen sequence and a MAP7 microtubule-binding domain (MTBD) fused to mNeonGreen via a GGGSGGGSTS linker (hereafter referred to as MAP7MTBD-mNeonGreen) by IDT Inc. Each fragment was inserted into the pJFRC81-10XUAS-IVS-Syn21-GFP-p10 backbone (Addgene 36432) using NotI (NEB catalog no. R3189) in combination with XbaI (NEB catalog no. R015S), respectively. For constructing the 1XUAS-IVS-Syn21-MAP7MTBD-mNeonGreen-p10 construct, the 1XUAS fragment was excised from pJFRC3-1XUAS-IVS-mCD8::GFP (Addgene 26216) using HindIII (NEB, catalog no.R3104S) and BglII (NEB, catalog no.R0144S), and inserted into the 10XUAS-IVS-Syn21-MAP7MTBD-mNeonGreen-p10 backbone. The 3XUAS-IVS-Syn21-MAP7MTBD-mNeonGreen-p10, 8XUAS-IVS-Syn21-MAP7MTBD-mNeonGreen-p10, and 15XUAS-IVS-Syn21-MAP7MTBD-mNeonGreen-p10 constructs were generated by inserting synthesized UAS fragments (3X, 8X, or 15X repeats) into the 10XUAS-IVS-Syn21-MAP7MTBD-mNeonGreen-p10 backbone cloned by GeneScript Inc. Plasmid constructs were sequence-verified and injected into *Drosophila* melanogaster embryos carrying the attP2 docking site for φC31 integrase–mediated germline transformation by BestGene Inc.

### Western blotting

Third-instar larvae at 120 h were collected for protein extraction. The control group is consisted of *w*; ppk-Gal4/Sp,Cyo; ppk-Gal4::UAS-EB1-RFP::10XUAS-MAP7MTBD–mNeonGreen::UAS-CD4IFP/Dr*, and the tubulin-reduced group consisted of *w*; ppk-Gal4/Sp,Cyo; ppk-Gal4::UAS-EB1-RFP::10XUAS-MAP7MTBD-*mNeonGreen*::UAS-CD4IFP/αTub84B*^*attp-KO*^. For each biological replicate, approximately 20 larval fillets were dissected in cold PBS. Tissues were transferred into pre-chilled RIPA buffer (Thermo scientific, catalog no. 89900) with protease inhibitors and thoroughly homogenized using a micro-pestle before lysis. Homogenates were incubated on ice for 30 min and centrifuged at 14,000 × g for 10 min at 4 °C to remove debris. Protein concentrations were measured using a Nanodrop. Equal amounts of total protein were mixed with Laemmli sample buffer and boiled at 95 °C for 10 min.

Proteins were resolved on 12% SDS–PAGE gels and transferred onto PVDF membranes. Membranes were blocked in 3% BSA in TBST for 1 h at room temperature, followed by overnight incubation at 4 °C with primary antibodies of mouse monoclonal anti–acetylated tubulin (Sigma Aldrich, catalog no. T6793) and rabbit monoclonal β-actin (GeneTex, catalog no. GTX637675). After washing, membranes were incubated with HRP-conjugated secondary antibodies for 1 h at room temperature.

Protein bands were detected using DAB substrate kit (Thermo scientific, catalog no. 34002) and imaged with a digital imaging system. Band intensities were quantified using ImageJ. Acetylated tubulin levels were normalized to β-actin, and statistical analyses were performed using Welch’s t-tests.

### Live Imaging of Class IV Neurons in *Drosophila* Larvae

All flies were maintained on apple juice agar plates with yeast paste at 25 °C and 60% humidity. Embryos were collected every 2 h and allowed to develop for ~120 h under the same conditions. Selected larvae were rinsed twice with DI water and anesthetized for 1.5 minutes by placing them next to tissue paper containing 1 mL of isoflurane (Covetrus) in a covered chamber. The larva was placed on a slide containing a 2% agar pad (~0.34 mm thick) with the dorsal side facing up and then covered with a coverslip. Imaging was performed on a spinning disk confocal microscope (Nikon CSU-W1) equipped with a Plan Apochromat 40× water-immersion objective (NA 1.25). Live imaging was performed using simultaneous laser excitation at 488 nm (30%) and 561 nm (35%) for 15 min, capturing 15-20 sections at 0.8 µm per section, with images collected at ~3.8 s intervals. After recording the movies, static images of the dendritic branches and surrounding subtree were acquired using 100% 640 nm laser excitation to capture the whole static neuronal morphology.

### Mesh size measurement

To quantify dendritic mesh size and morphological changes, we adapted the method described by Ganguly et al. (86). Briefly, dendritic arbors were first skeletonized from confocal membrane images. To estimate the characteristic mesh size, we calculated the *hitting probability* H(B), defined as the probability that a randomly placed box of side length *B* intersects the dendritic skeleton. The probability *H*(*B*) increases monotonically toward 1 as *B* approaches the total size of the neuron. The characteristic mesh size *B*_*H*_ was defined as the box size at which half of all boxes hit the dendrite. All computations were implemented in custom MATLAB scripts.

### Microtubule dynamic parameter measurements

Four dynamic parameters: polymerization rate *v*_*g*_, depolymerization rate *v*_*s*_, catastrophe frequency *f*_*c*_, and rescue frequency *f*_*r*_ were quantified from kymographs. To characterize microtubule catastrophe, all full growth events were analyzed. Events that were spatially cut off (growing out of the field of view) and temporally cut off (starting to grow prior to the onset of the time-lapse or still growing at the end of the time-lapse) were disregarded. Rescue frequency was reported as the ratio of the total number of rescues observed over the total amount of time microtubules spent during depolymerization. Standard deviations for catastrophe frequency and rescue frequency were calculated as 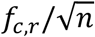 (assuming a Poisson distribution), where *n* is the number of counted events.

The relationship between the microtubule four dynamic parameters and leaf number was calculated as follows. Growth and shrinkage velocities were assigned to every leaf number that a microtubule passed through during the corresponding growth or shrinkage segment. In contrast, catastrophe and rescue frequencies were calculated using only the leaf number at which each respective event occurred, corresponding to the end point of a growth or shrinkage phase.

Because the neuronal arbor is a branched structure, leaf numbers are densely populated at small leaf numbers and sparsely populated at large ones. To account for this non-uniform distribution, we adopted logarithmic binning to the data. Only logarithmic bins with contributions from at least three neurons, each providing a minimum of six events in that bin, were considered.

### Branch intensity measurements

For each centerline point at position with coordinates (*x*_*c*_, *y*_*c*_), a local normal vector was computed from its neighboring two centerline points. Intensities were then sampled along this normal within a ±11 px window centered at (*x*_*c*_, *y*_*c*_). The position *x*_*n*_ of the peak intensity along the normal was identified, and the signal was integrated within a ±5 px window around this peak. This process was repeated for every centerline point, yielding a one-dimensional intensity profile along each branch denoted by *I*(*x*_*c*_, *t*).

### Fluorescence Recovery After Photobleaching (FRAP)

We performed two sets of FRAP experiments: the first one measured the diffusion coefficient of cytoplasmic mNeonGreen fluorophores, the second one quantified the contrast of MAP7MTBD–mNeonGreen.

A small rectangular region along a single dendritic branch was photobleached with a 405 nm laser (power: 40%, dwell time: 20 µs). For the first FRAP experiment, *w*; ppk-gal4/10XUAS-mNeonGreen;ppkCD4tdTomato/Dr* was used. After photobleaching, time-lapse imaging was performed before and after bleaching at 40 frames per second (25 ms exposure). For the second FRAP experiment, *w*;ppk-Gal4/Sp,Cyo;ppk-Gal4::UAS-EB1-RFP::10XUAS-MAP7MTBD-mNeonGreen::UAS-CD4-IFP/Dr* flies were used. Time-lapse imaging was performed at 10 frames per second (100 ms exposure) after photobleaching.

### Preprocessing for FRAP data

Double normalization involves the normalization of the recovery signal to the average prebleach signal and, at the same time, considers the loss of total signal due to the bleaching during the postbleach imaging. For double normalization, the average intensity of fluorescent signal in three regions of interest is measured for each time point: the bleached region *I*(*t*), the unbleached region *U*(*t*), and a region outside the cell to determine the background signal (BG). BG was estimated for each experiment as the mean pixel intensity measured over visually identified background regions located far from any dendrite, across 10 representative movies. Here are the four steps: 1. background subtraction: calculate for each time point *t, I*(*t*) − *BG* and *U*(*t*) − *BG*; 2. determine the lost fluorescence signal: calculate for each time point *t*, [*U*_*prebleach*_ − *BG*]/[*U*(*t*) − *BG*]; 3. correct for lost signal due to bleaching during imaging: calculate for each time point *t*, [*I*(*t*) − *BG*] [*U*_*prebleach*_ − *BG*]⁄[*U*(*t*) − *BG*]; 4. determination of the relative fluorescence signal in the bleached region: calculate for each time point *t*:

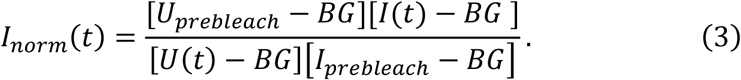

### Method to determine the size of the FRAP area and diffusion coefficient

The evolution of the protein concentration of the unbleached fluorescent molecules *φ*(***x***, *t*) during the FRAP experiment is given by:

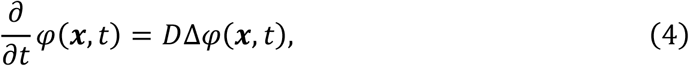

where *D* is the diffusion constant. Here we only consider the one-dimensional case in which *φ* varies with *x*.

Taking the Fourier transform w.r.t. the spatial variables we have:

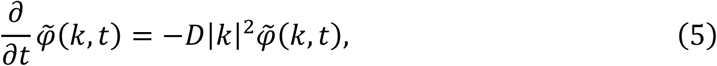

with initial condition 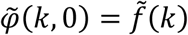. This equation has a unique solution satisfying the initial conditions, given by: 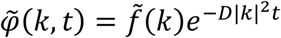. The convolution theorem tells us that solution *φ*(*x, t*) will be the convolution of the initial condition *f*(*x*) with the inverse Fourier transform of the Gaussian. We have 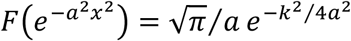. Therefore, 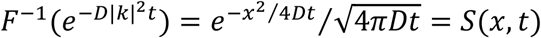.

Putting everything together, the general solution to Eq. (4) is:

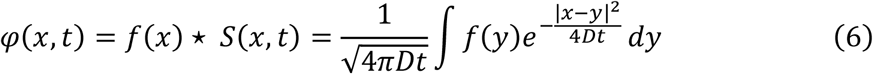

The initial condition of *f*(*x*) takes the form of:

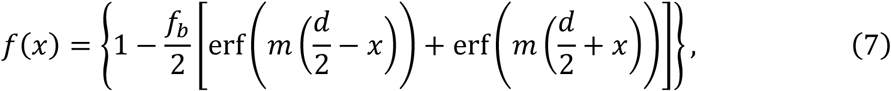

here *d* stands the extent of the bleached region, the steepness of the boundaries is described by *m*. The fraction bleached is specified by *f*_*b*_. We can get these parameters from the first postbleach image.

The time evolution of the fluorescent protein density is given by:

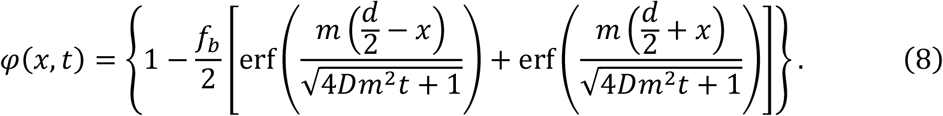

The mean fluorescence within the bleached region is given by:

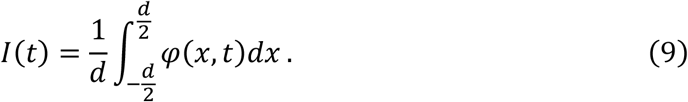

The normalized fluorescence recovery curve is given by:

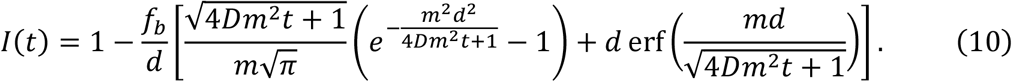

When 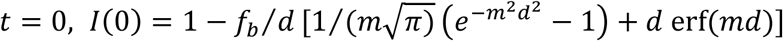, when *t* → ∞, *I*(*t*) → 1.

When there exists asymmetry of the two sides of the bleached region, the initial condition of *f*(*x*) takes the form of:

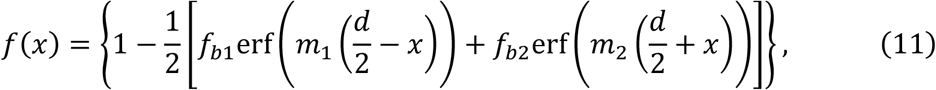

where 1 and 2 stand for the two sides.

The corresponding one-dimensional intensity profile along the branch reveals partial bleaching at the FRAP region boundaries and is fitted with Eq. (11), yielding the parameters *f*_*b*1_, *f*_*b*2_, *m*_1_, *m*_2_ and FRAP region width *d*.

The time evolution of fluorescent intensity is given by:

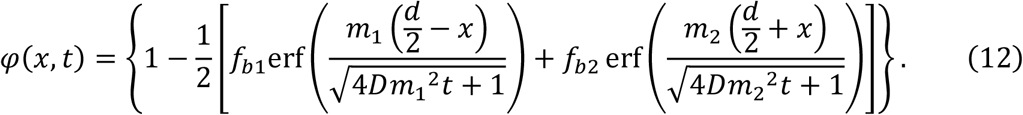

The mean fluorescence within the bleached region is:

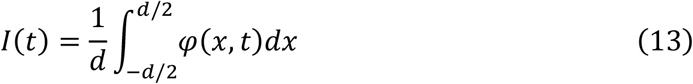

Further normalization leads to:

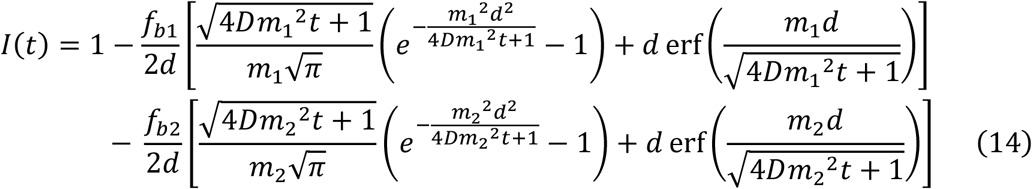

The diffusion coefficient of mNeonGreen fluorophores was obtained by fitting Eq. (14) using the fitted parameters from Eq. (11). We observed no significant difference between the measured diffusion coefficients in the primary and peripheral branches, and measured free mNeonGreen diffusion coefficient of *D*_mNG_ = 32.0 ± 15.9 μm^2^ ⋅ s^−1^ (mean ±SD, data points from six animals). In water (viscosity 1cP), the measured diffusion coefficient for GFP is 87 μm^2^ ⋅ s^−1^(87). Thus, the effective viscosity in dendrites is ~2.7cP.

To estimate the diffusion coefficient of cytoplasmic tubulin, we applied the Stokes-Einstein relation and used the molecular weights of mNeonGreen (~26.6 kDa) and a tubulin dimer (~110 kDa). The estimated diffusion coefficient is *D*_αβ−tub_ = 20 ± 10 μm^2^ ⋅ s^−1^.

### Single microtubule intensity measurement

Single microtubule intensity was measured from individual MT events, either growth or shrinkage events, identified in low leaf number branches. For each event, we manually annotated the spatial coordinates and the corresponding time points marking the start and end of shrinkage. The fluorescence intensity at the start includes both cytoplasmic and MAP7MTBDmNeonGreen signal *I*_cyto_ + *I*_MT_, whereas the intensity at the end represents cytoplasmic signal *I*_cyto_.

Since mNeonGreen photobleaching is negligeable during a dynamic event (typically ~100 s), we computed single MT intensity taking the difference between the two time points. For each neuron, we measured ~10 single MT intensities at different locations. Mean and standard deviation of single MT intensities was finally seeded to our intensity profile analysis pipeline.

### Statistics and reproducibility

Statistical analyses were performed in Excel, GraphPad Prism and MATLAB (Version R2024a). Variables were analyzed using Welch’s t tests for two groups of data, and Brown-Forsythe ANOVA test for data with more than two groups. Reproducibility was confirmed by at least three independent experiments.

## Acknowledgments

We are grateful for helpful discussions and support from colleagues and friends. We thank Jill Wildonger for providing tubulin knockout flies, and Jeffrey Fredberg, Howard Stone, and Dan Needleman for valuable feedback on the manuscript. We also thank Joe Howard, Xiao Xiao, Yuhai Tu, Mark Williams, Paul Champion, Armen Stepanyants, and Ping Sheng for valuable discussions at different stages of the project. We thank Yi Lin and Xuan Guo for their assistance with fly injections.

## Funding

Northeastern University Mechanobiology Postdoctoral Fellowship (L.T.) NIH R01 GM072744 (N.W.)

NIH R35 GM131753 (T.J.M.)

CASI Burroughs Wellcome Fund (M.L.)

## Author Contributions

D. Y. and M. L. performed experiments; L.T., M.L. and D. Y. conducted the data analysis; L.T. and M. L. performed the theoretical modeling and simulations; Z. X. and Y. F. developed the machine learning pipeline; M.L., T.J.M., N. W., and Y. F. supervised different aspects of the project; M.L conceived the project, oversaw its implementation, and wrote the manuscript. All authors contributed to manuscript editing.

